# Evaluation of training data, ground truth and shape variability in the semantic segmentation of HeLa cells observed with electron microscopy

**DOI:** 10.1101/2022.06.01.494316

**Authors:** C. Karabağ, C.C. Reyes-Aldasoro

## Abstract

This paper describes the quantitative and qualitative evaluation of training data, ground truth and the variability of the shape of HeLa cells as observed with electron microscopy in the process of semantic segmentation. HeLa cells are a widely used immortal cell line, but the principles described here could apply to other cells. From a data set of 8, 000*×*8, 000*×*517 voxels, a region of interest (ROI) of 2, 000*×*2, 000*×*300 was cropped for quantitative evaluation with the qualitative evaluation performed on a larger section of interest. Patches of data and labels were used to train a U-Net architecture from scratch, which was compared also with a traditional image processing algorithm. The correctness of ground truth, that is, the inclusion of one or more nuclei within a region of interest was evaluated with a hand-drawn ground truth of the ROI. The impact of the extent of training data was evaluated by comparing results from 36,000 data and label patches extracted from the odd slices in the central region, to 135,000 patches obtained from every other slice in the set. Accuracy and Jaccard similarity index were higher for the larger training set with multiple nuclei in the ground truth. However, when the U-Nets were used to segment the section of interest beyond the ROI, results were not as good as expected, suggesting that the very complicated nature of the shapes of HeLa cells cannot be learned from a single cell. Data, code and ground truth have been made publicly released through Empair, GitHub and Zenodo.

## 1 Introduction

The immortal *HeLa* cell line, which originated from cervical cancer cells of the patient Henrietta Lacks, is the oldest and most commonly used human cell line [14, 18]. These cells have been widely used in numerous scientific experiments, and have recently became subject of books and films based on the life of Henrietta Lacks, the extraction of the cells, which were removed, stored and distributed without the consent or knowledge of the patient or her family, as this was not a requirement in 1951 in the United States, and the subsequent story that the cells followed. The remarkable and engaging story is narrated by Rebecca Skloot in her book *The immortal life of Henrietta Lacks* [21] and Henrietta Lacks has been portrayed by Oprah Winfrey in a film with the same name.

One area of interest in cancer research is the observation of the distribution, shape and morphological characteristics of the cells and cellular structures like the nuclear envelope and the plasma membrane [19, 23, 24, 26]. Whilst it is possible to observe these characteristics of the cells visually, or to manually delineate the structures of interest [1, 17, 27, 2], either by a single expert or through citizen-science efforts [8], automatic segmentation of the cell and its structures is crucial for high throughput analysis where large amounts of data, i.e. terabytes [9], is acquired regularly.

The segmentation and classification of cells and their structures remains as one important and challenging problem of interest for the clinical and programming communities [4, 25]. Segmentation nuclear envelope (NE) and the plasma membrane depends on the resolution of the acquisition equipment and the contrast it provides, as well complexity of the structures themselves. At higher resolutions and in three dimensions, such as those provided by electron microscopy, the problem is challenging [15, 16]. Segmentation with traditional image processing algorithms and deep learning approaches [13] are widely used in tasks of segmentation, and have previously been compared for the segmentation of nuclear envelope and plasma membranes of HeLa cells as observed with electron microscopes [7, 10, 11, 22]. In this work, the impact that the training data can have on the outcome of a segmentation was evaluated, and a comparison of the segmentations of HeLa plasma membranes and nuclear envelope with a U-Net [20] was performed. Specifically, a Region of Interest (ROI) of 2, 000 × 2, 000 × 300 voxels was cropped from a larger data set of 8, 000 × 8, 000 × 517 voxels, and was manually delineated to have a ground truth. The ROI was segmented under the following scenarios: (a) training a U-Net from scratch with 36,000 patches for data and labels from alternate slices of the central region of the cell (101 : 2 : 180) considering that there was a single cell in the ROI, (b) training a U-Net from scratch considering the existence of multiple cells in the ROI and 36,000 patches for data and labels from alternate slices of the central region of the cell (101 : 2 : 180), (c) training a U-Net from scratch considering a multiple cells in the ROI and 135,000 patches for data and labels from alternate slices of the whole cell (1 : 2 : 300). The results were compared slice-pre-slice against a previously published traditional image processing algorithm using accuracy and Jaccard similarity index. Further, the trained U-Nets were applied to segment the 8, 000 8, 000 slices from which the ROI had been extracted and the results were visually assessed as there was no ground truth available. All the programming was performed in Matlab^®^ (The Mathworks^™^, Natick, USA).

## 2 Materials and Methods

### 2.1 HeLa cells preparation and acquisition

Details regarding the preparation of the HeLa cells have been previously described [11]. For completeness these are briefly described. Cells were embedded in Durcupan and observed with an SBF Scanning Electron Microscope following the National Centre for Microscopy and Imaging Research (NCMIR) protocol. [3]. The images were acquired with a SBF SEM 3View2XP microscope (Gatan, Pleasanton, CA, USA) attached to a Sigma VP SEM (Zeiss, Cambridge, UK). Voxel dimensions were 10×10×50 nm with intensity [0–255].

Five hundred and seventeen images of 8, 000 × 8, 000 pixels were acquired, from which a ROI was cropped by estimating manually the centroid of the cell to centre a box of of 2, 000 × 2, 000 × 300 voxels. Fig. 1 illustrates a representative slice of one of the 8, 000 × 8, 000 images. Fig. 2 illustrates several slices of the 2, 000 × 2, 000 ROI. EM data are publicly available through EMPIAR [5] http://dx.doi.org/10.6019/EMPIAR-10094).

**Fig. 1.**
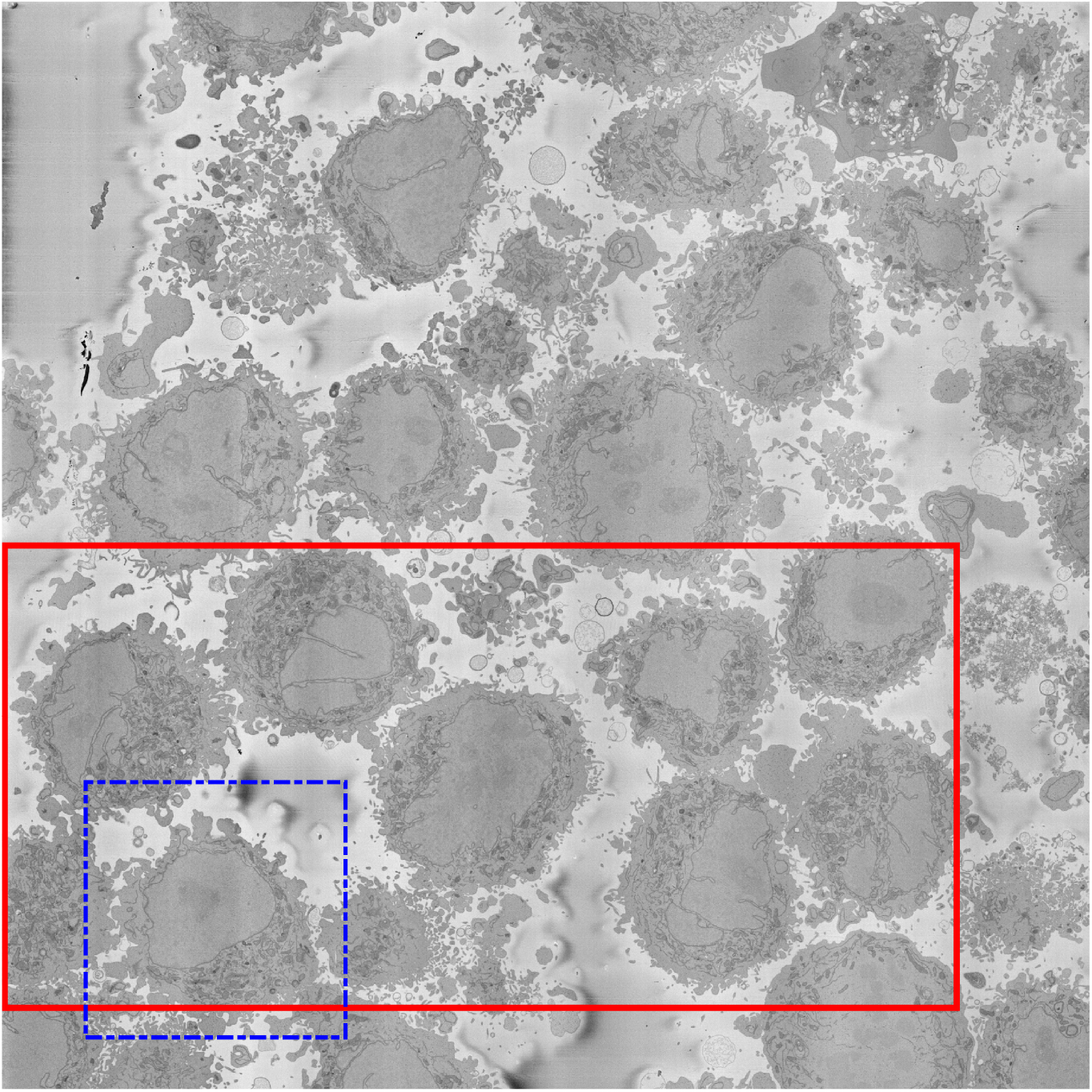
One representative slice of one of the 8, 000 *×* 8, 000 *×* 517 images. Several HeLa cells with different characteristics are visible. A solid red box shows an area of interest for future analysis and one cell that was selected as a region of interest (ROI) is at the bottom left surrounded with a dashed blue box. Several HeLa cells with different characteristics are visible. Notice how the nucleus of the cell of the ROI is relatively smooth and regular as opposed to other nuclei that present invaginations and more complex structures.

**Fig. 2.**
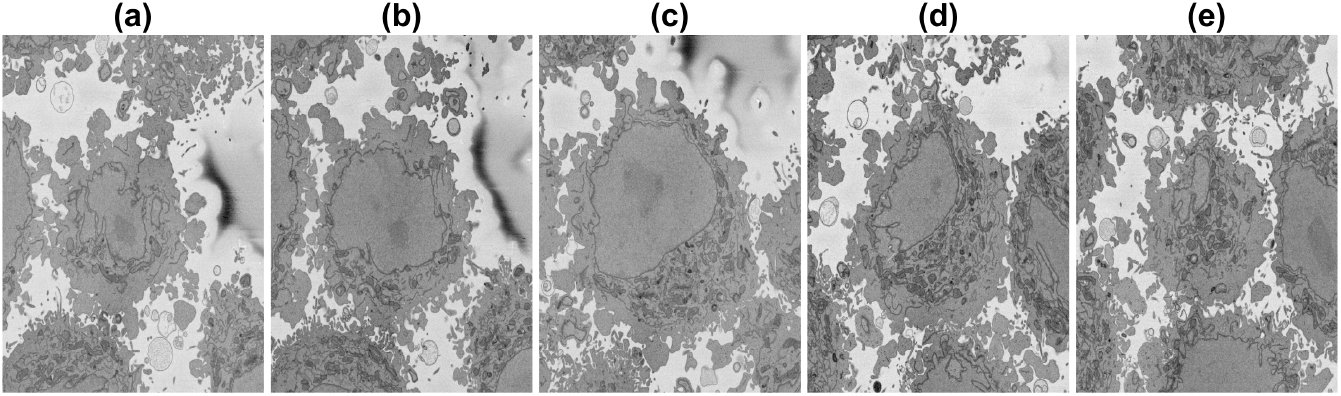
Five representative slices of the 2, 000*×*2, 000*×*300 region of interest (ROI) with the central slice (c) corresponding to the same image as Fig. 1. It should be noticed the cells that surround the central cell, nuclei are particularly visible towards the top (e) at the right and bottom of the image.

### 2.2 Ground truth (GT)

Ground truth for the 2, 000 × 2, 000 × 300 ROI was manually delineated sliceper-slice in Matlab. Disjoint nuclear regions were assessed by scrolling up and down to decide to include as part of the nucleus. The ground truth for ROI with only the central nucleus and the central nucleus and other nuclei are publicly available through Zenodo: https://doi.org/10.5281/zenodo.3874949 for single nucleus, https://doi.org/10.5281/zenodo.6355622 for multinuclei.

### 2.3 U-Net training data, architecture and segmentation

The U-Net architecture [20] is convolutional neural network architecture, which has been widely used for semantic segmentation among other tasks. The essence of the U-Net architecture is a combination of downsampling steps obtained by convolutions and downsampling, which are followed by upsampling steps through which the classification obtained at lower resolutions is propagated upwards towards higher resolutions with a rough shape of a letter “U”, hence it is called U-Net. The architecture can be trained end-to-end from pairs or patches of images and their corresponding classes.

For this work, a semantic segmentation pipeline with a U-Net architecture was implemented in Matlab. Three different scenarios were followed in the preparation of the training data. First, patches of images and labels of size 128 × 128 with 50% overlap were generated from 40 alternate slices of the central region of the cell (101 : 2 : 180) considering that there was a single cell in the ROI. For one 2, 000 × 2, 000 image, there were 30 × 30 patches and thus 30 × 30 40 corresponded to 36, 000 patches. Alternate slices were selected to exploit the similarity between neighbouring slices and only the central region was selected to provide a reduced set, which would hopefully capture the variability of the cell.

In this case, the ground truth included only the nucleus of the central cell visible in the ROI. Second, the same strategy was followed to generate 36,000 patches and labels from alternate slices of the central region of the cell (101 : 2 : 180), however, in this case, the ground truth included the nuclei of all cells visible in the ROI. Finally, the ground truth was extended to 135,000 patches for data and labels from alternate slices of the whole ROI (1 : 2 : 300), in this case, there were 150 slices and 900 patches per slice.

The U-Net architecture was tuned during 15 epochs, Adam optimiser [12], input images of 128 *×* 128, initial learning rate of 1e-3, a mini batch size of 64, and configuration shown in Fig. 3. Once the network was trained, it was used to segment the 300 images of the ROI and the original 8, 000 × 8, 000 images. Finally, a simple post-processing consisting of the following steps was applied: filling holes, closing with a disk structural element of 3 pixel radius, removal of small regions (area *<* 3,200 pixels, equivalent to 0.08% of the area of the image). Matlab code is available through GitHub: (https://github.com/reyesaldasoro/HeLa_Segmentation_UNET2).

**Fig. 3.**
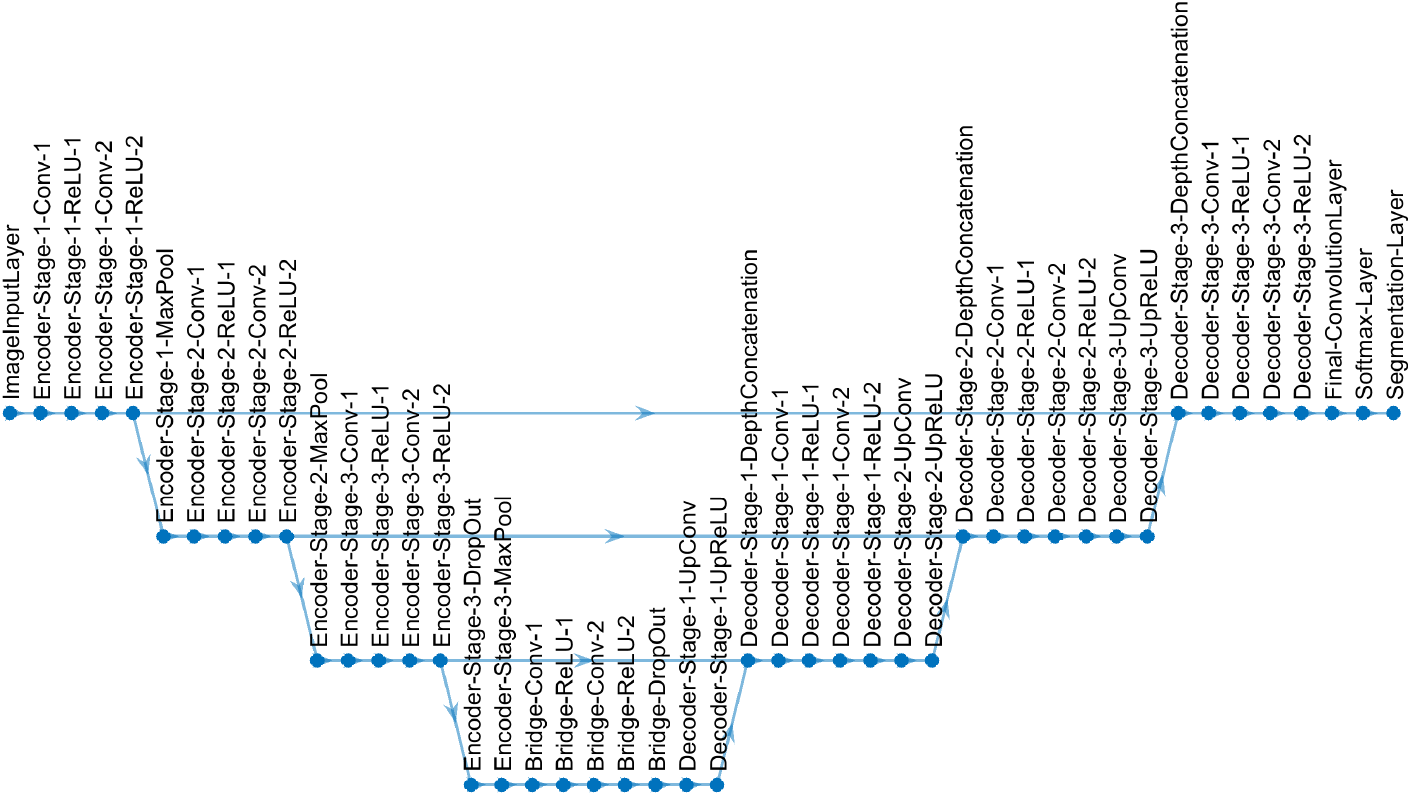
U-Net architecture used for semantic segmentation in this work.

### 2.4 Image analysis segmentation algorithm

The semantic segmentations obtained with the different configurations of the U-Net were compared against a previously published algorithm [9] based on traditional image analysis steps: low pass filtering, edge detection, generation of superpixels, morphological operators and selection by size and location, combination of adjacent slices to select disjoint regions that belong to the nucleus. This algorithm obtained accurate semantic segmentations which outperformed other deep learning approaches [10]. However, in the case of U-Net, it was noticed that the inferior accuracy and Jaccard index were due to the discrepancy between the segmentation, which detected several nuclei, and the ground truth, which considered only the central nucleus. (link removed).

### 2.5 Quantitative comparisons

The segmentations assigned a class (background, nucleus, nuclear envelope, rest of the cell) to each pixel of the images. The accuracy and Jaccard similarity index (JI) [6] were calculated pixel-by-pixel on a comparison of the ground truth and results for the nuclei. Accuracy corresponded to (*TP* + *TN*)*/*(*TP* + *TN* + *FP* + *FN*) and JI, or intersection over union of nuclear area corresponded to (*TP*)*/*(*TP* + *FP* + *FN*), where TP corresponds to True Positives, TN to True Negatives, FP to False Positives and FN to False Negatives.

## 3 Results and Discussion

Segmentation of all slices and calculation of accuracy and Jaccard index was performed as described in previous sections. Figs. 4, 5, 6 illustrate the results obtained with the image processing algorithm and the U-Net with different training conditions.

**Fig. 4.**
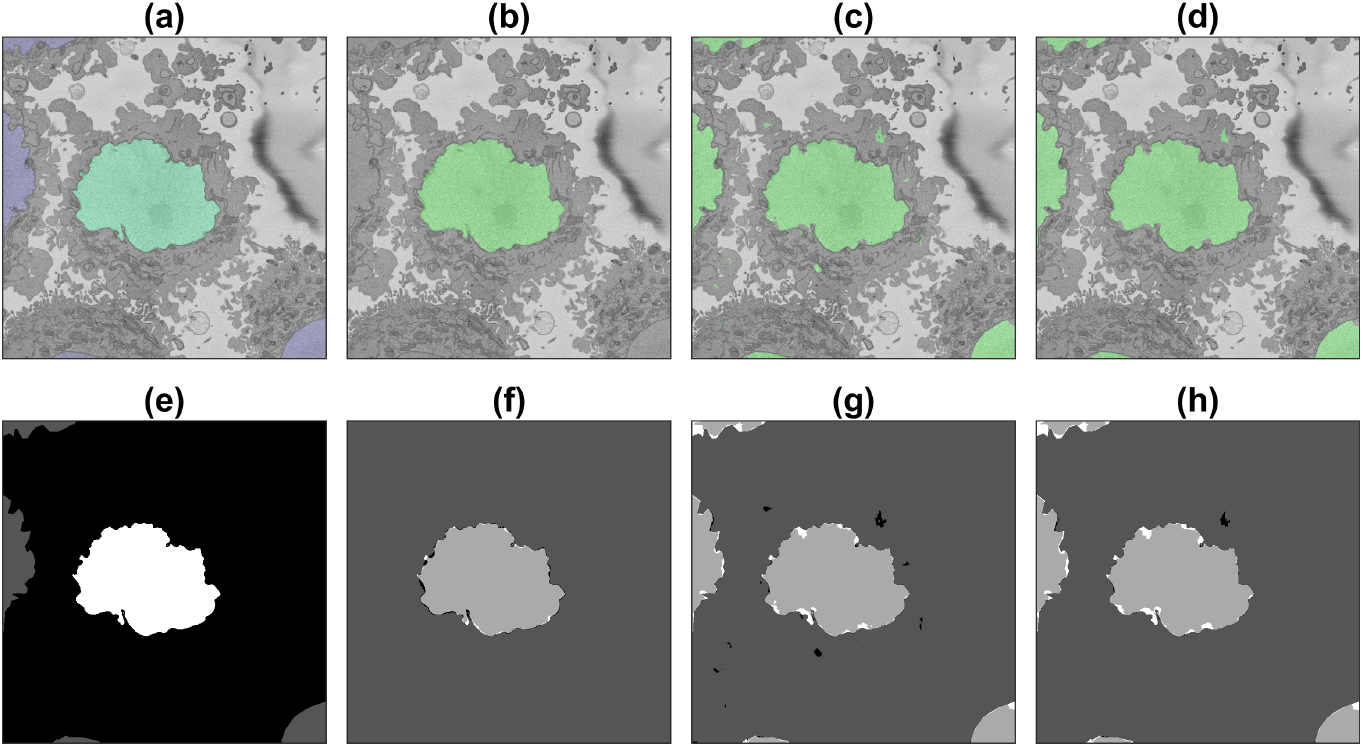
Illustration of the semantic segmentation of one slice (71/300) of the ROI with the image processing algorithm and U-Net (36,000 patches). Top row (a-d) show slice with results overlaid, bottom row (e-h) shows ground truth (GT) and results as classes. (a,e) GT, central nucleus in green/white whilst other nuclei in purple/gray and background in grayscale/black. (b,f) IP segmentation. (c,g) U-Net segmentation. (d,h) U-Net segmentation after post-processing. In (f-h) True positives = light gray, True negatives = darker gray, False positives = black, False Negatives = white.

**Fig. 5.**
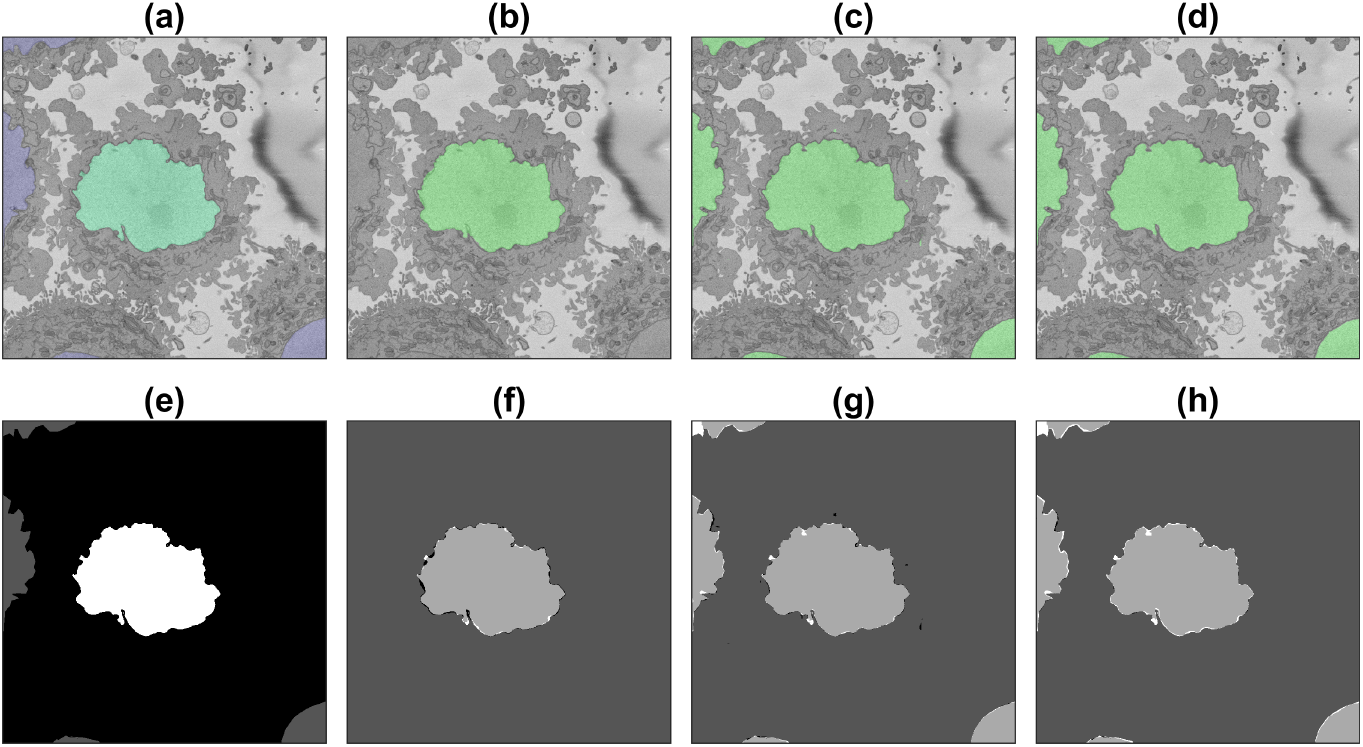
Illustration of the semantic segmentation of one slice (71/300) of the region of interest (ROI) with the image processing (IP) algorithm and U-Net trained with 135,000 patches. For a description, see Fig. 4.

**Fig. 6.**
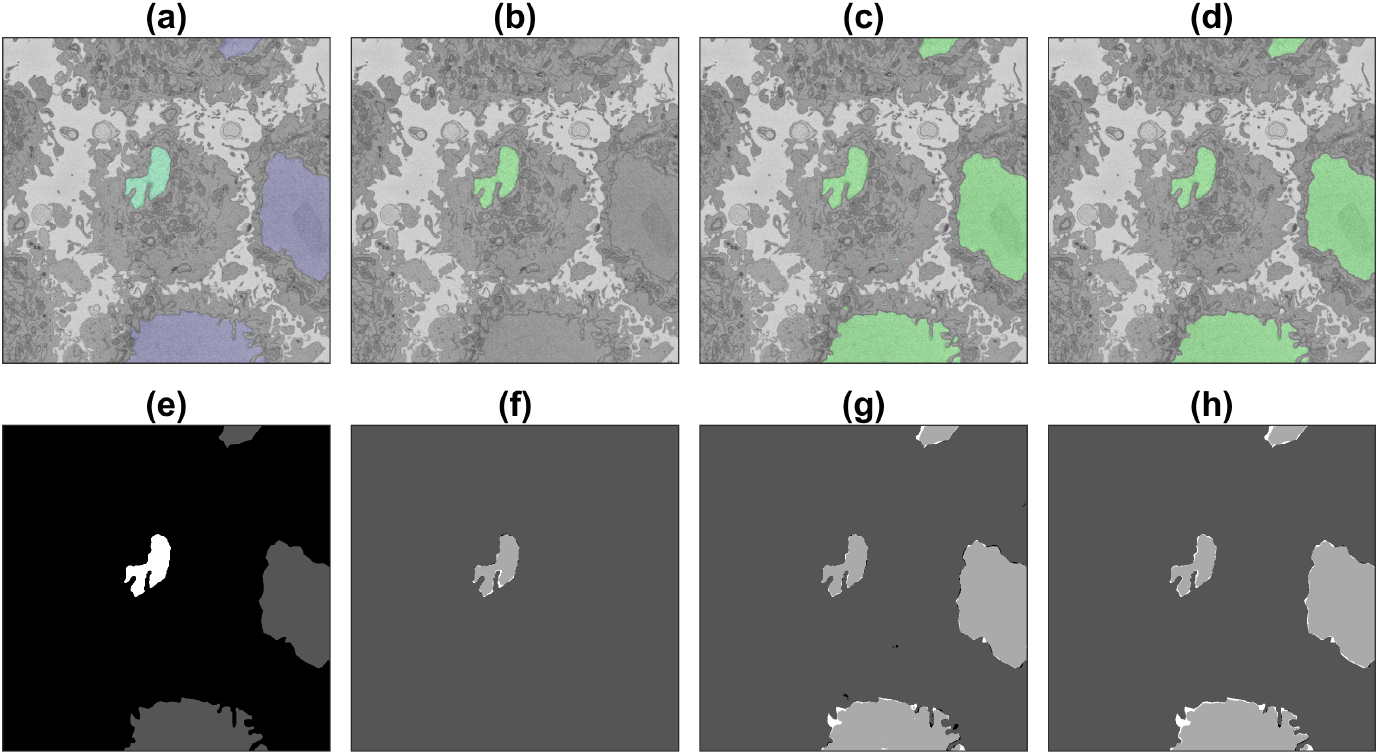
Illustration of the semantic segmentation of one slice (251/300) of the region of interest (ROI) with the image processing (IP) algorithm and U-Net trained with 135,000 patches. For a description, see Fig. 4.

Accuracy and Jaccard index results are shown in Fig. 7. Several points are worth mentioning. When the U-Net was trained and measured with GT with a single nucleus (blue line in Fig. 7), the results were good between slices 150 and 200 of the ROI as these slices contained only one nucleus. Towards the top and bottom of the ROI the results dropped as other nuclei appeared in the images, and these were segmented, but were not part of the GT and consequently numbers dropped. The values of the Jaccard index behaved in a similar way, high in the central slices and low on the top/bottom slices. The results of the image processing algorithm (thin dashed black line in Fig. 7), which segmented only the the central cell, and were compared against the GT that contained only the central cell produced very good results, especially towards the central region of the cell, where the nucleus is largest. Towards the top and bottom of the cell, where the nucleus is smaller and the shape less regular, the accuracy remained high due to the large number of TN, but the JI dropped in a similar way to the U-Net with a single nucleus and was zero in the extremes due to absence of TP.

**Fig. 7.**
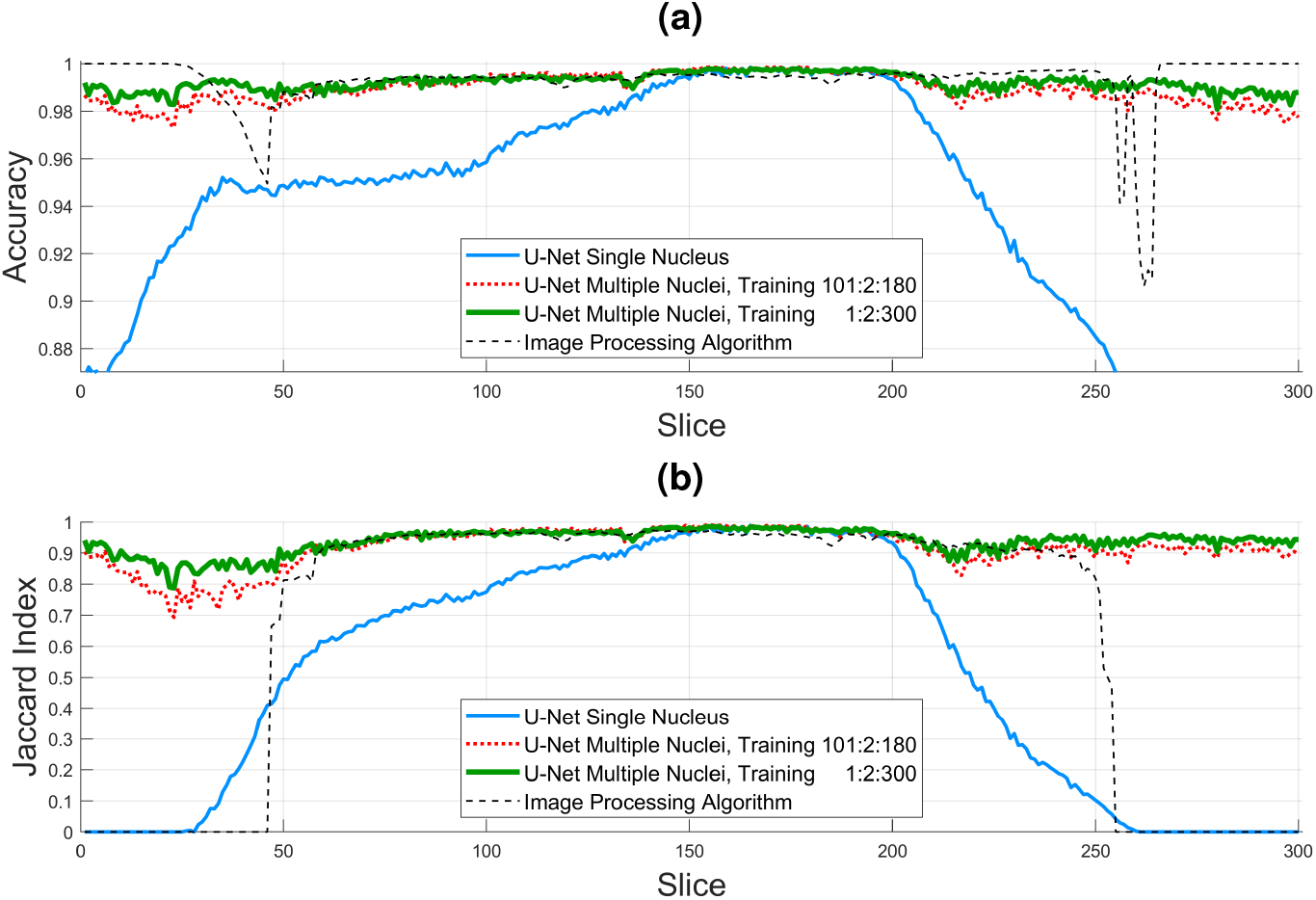
Numerical comparison of the four algorithms (U-Net single nucleus, U-Net multiple nuclei 36,000 patches, U-Net multiple nuclei, 135,000 patches, image processing (IP) algorithm). (a) Accuracy. Average values for all slices (0.9346, 0.9895, 0.9922, 0.9926), for slices 150:200 (0.9966, 0.9974, 0.9971, 0.9945). (b) Jaccard Similarity Index (JI). Average values for all slices (0.5138 0.9158 0.9378 0.6436), for slices 150:200 (0.9712, 0.9778, 0.9760, 0.9564).

For the U-Net trained with multiple nuclei and 36,000 patches (dotted red line in Fig. 7) the results were significantly higher than the single nucleus UNet, close to the image processing algorithm in central slices and better in the top/bottom as both metrics remained high. However, for the U-Net with 135,000 patches (thick green solid line in Fig. 7) the results were higher still, especially in the regions outside the slices from which the 36,000 patches were generated (i.e. 1 : 2 : 99 and 181 : 2 : 300). The average accuracy for all slices for the four algorithms were: U-Net single nucleus, **0.9346**, U-Net multiple nuclei 36,000 patches, **0.9895**, U-Net multiple nuclei, 135,000 patches, **0.9922**, image processing algorithm **0.9926**, which in general are very high for all cases where the appropriate GT is considered, but the high values also indicate a large number of TN, especially on the top and bottom slices. For the central slices (150 : 200) where there is a single cell and results are best the values are all above 99% (same order **0.9966, 0.9974, 0.9971, 0.9945**), thus the errors are due to small variations on the edge of the nucleus or the fact that the manual delineation can also include small errors. The values for the Jaccard index are more interesting, with the average values for all slices of (**0.5138 0.9158 0.9378 0.6436**), it is clear that the multinuclei U-Nets have a significant improvement as they consider better the top and bottom slices. For the central slices 150 : 200 the values are pretty similar for the three U-Nets and slightly lower for the image processing (**0.9712, 0.9778, 0.9760, 0.9564**) as a small dip is visible around slice 180. A final comparison in Jaccard, between slices 60 : 150, where the curves are very similar shows the following values (**0.8047, 0.9579, 0.9592, 0.9565**), that is, nearly identical averages for all methods except the initial U-Net. These results confirm that the larger the training data, the better the U-Net can learn the characteristics of the cells. It is important to highlight that the results are not completely like-with-like as the GT is not the same for the single nucleus and the multinuclei approaches.

As a further segmentation test, the two U-Nets trained on multiple nuclei were used to segment the 8, 000 × 8, 000 slices and the results are shown in Fig. 8 together with the image processing algorithm. The first observation is that the U-Net with the larger training set (b) provides better results than the one with smaller training set (a). This can be observed in several locations, like the small notch of the cell towards the bottom left, the background classified as cell in the dark region just above the same cell, or the nucleus of the cell towards the top right. A second important observation is that not all nuclei are well segmented, as those on the top right. These can be compared with the image processing algorithm (c) which provide better results for those two cells. These less than satisfactory results may be the outcome of training the U-Nets on a a single cell, whose nucleus is considerably smoother than those of the surrounding cells. It can be speculated that a larger number of training patches, *from different cells*, may be necessary to obtain better results.

**Fig. 8.**
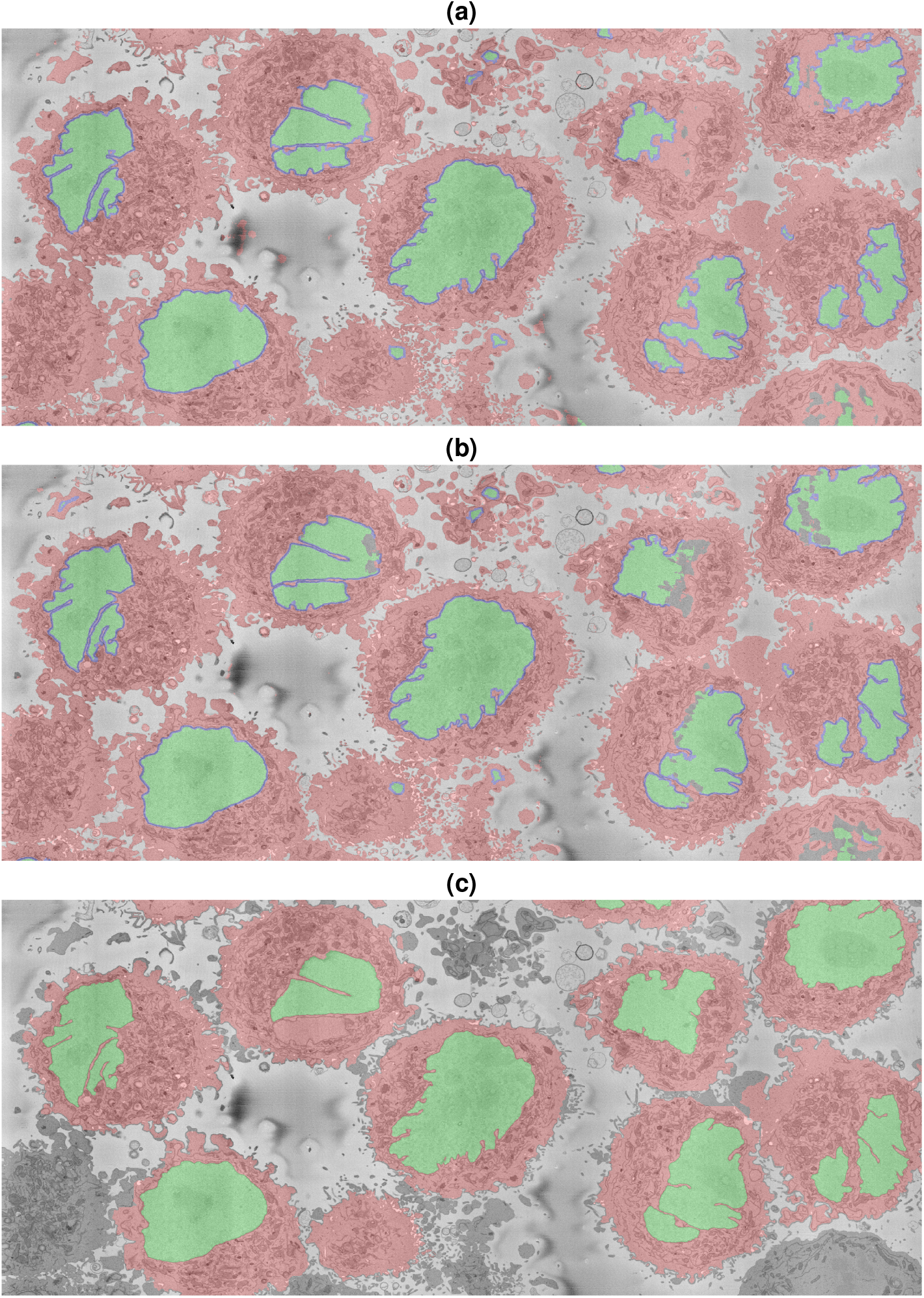
Segmentation of the section of interest shown in Fig. 1 (red box). (a) U-Net trained with 36,000 patches. (b) U-Net trained with 135,000 patches. (c) Image processing (IP) algorithm. Cells are shown with a red shading, nuclei are shown with a green shade and for the U-Net results, the nuclear envelope is shown with a blue shade.

The image processing algorithm was run on an instance segmentation and only a number of cells were selected, which can be noticed in that some cells (bottom right and left) are not segmented and appear gray. It can also be noticed that some cellular regions are not segmented, especially those that are between cells. However, the shapes of the nuclei appear better segmented, with the exception of the second cell from the left on the top, for which a section of the nuclei was not detected.

## 4 Conclusion

The quantitative and qualitative comparison of different semantic segmentation approaches of HeLa cells, their nucleus, nuclear envelope, cell and background, have been presented in this work. Whilst, as would be expected, the fact that a larger amount of training data provides more accurate segmentation was sustained, it was further observed that the variability of the cells, even when it is a single line of cells, precludes the generalisation that training data can be extended from one cell to another without degradation of the results.

Furthermore, the consideration of the correct ground truth can be extremely important in the assessment of results of different approaches. The algorithms, data and ground truths have been released open source for the community.

## Acknowledgements

The authors acknowledge the useful discussions with Martin L. Jones and Lucy M. Collinson from The Francis Crick Institute, UK.

